# Impact of light on task-evoked pupil responses during cognitive tasks

**DOI:** 10.1101/2023.04.12.536570

**Authors:** Islay Campbell, Elise Beckers, Roya Sharifpour, Alexandre Berger, Ilenia Paparella, Jose Fermin Balda Aizpurua, Ekaterina Koshmanova, Nasrin Mortazavi, Siya Sherif, Gilles Vandewalle

## Abstract

Light has many non-image-forming functions including modulation of pupil size and stimulation of alertness and cognition. Part of these non-image-forming effects may be mediated by the brainstem locus coeruleus. The processing of sensory inputs can be associated with a transient pupil dilation that is likely driven in part by the phasic activity of the locus coeruleus. Here, we aimed to characterise the task-evoked pupil response associated with auditory inputs under different light levels and across two cognitive tasks. We continuously monitored the pupil of 20 young healthy participants (24.05y ±4.0; 14 women) while they completed an attentional and an emotional auditory task whilst exposed to repeated 30-to-40s-blocks of light interleaved with darkness periods. Blocks could either consist of monochromatic orange light [0.16 melanopic Equivalent Daylight Illuminance (EDI) lux] or blue-enriched white light of three different levels [37, 92, 190 melanopic EDI lux; 6500K]. For the analysis 15 and then 14 participants were included in the attentional and emotional tasks respectively. Generalized Linear Mixed Models showed a significant main effect of light level on the task-evoked pupil responses triggered by the attentional and emotional tasks (*p*≤.0001). The impact of light was different for the target vs. non-target stimulus of the attentional task but was not different for the emotional and neutral stimulus of the emotional task. Despite a smaller sustained pupil size during brighter light blocks, a higher light level triggers a stronger task-evoked pupil response to auditory stimulation, presumably through the recruitment of the locus coeruleus.

## Introduction

The non-image-forming (NIF) system (also termed non-visual system) in the human retina detects environmental irradiance to mediate the influences of light on many NIF functions, including circadian entrainment (Berson et al., 2002), melatonin suppression (Brainard et al., 2001), pupillary light responses (Gamlin et al., 2007; Hattar et al., 2002), and stimulation of alertness and cognitive performance (Vandewalle et al., 2009). The primary photoreceptors of this system are intrinsically photosensitive retinal ganglion cells (ipRGCs) (Mure, 2021; Provencio et al., 2000), which express the photopigment melanopsin. Animal studies have established that the ipRGCs project to various subcortical brain regions, including the suprachiasmatic nucleus (SCN) of the hypothalamus, the site of the master circadian clock (Tri & Do, 2019). The exact brain pathways involved in light’s NIF functions for humans is an area of continued and active research. The locus coeruleus (LC), in the brainstem, receives indirect inputs from the SCN, and it is hypothesised that the LC may be involved in mediating light’s influence on alertness and cognition (Aston-Jones et al., 2001; Aston-Jones & Cohen, 2005; Vandewalle et al., 2009). The LC is central to cognition and alertness and a major source of norepinephrine (NE) in the brain (Aston-Jones & Cohen, 2005). Neuroimaging studies reported that an area of the brainstem compatible with the LC is modulated by the wavelength of light while performing a non-visual cognitive task (Vandewalle et al., 2007, 2009). The LC has a deep location in the brain stem and is small in size, approximately 0.15mm long and 2.5mm in diameter (∼50.000 neurons in total) (Keren et al., 2009). Therefore, its role in mediating the NIF impacts of light is difficult to assess. Here, we emphasize that variation in pupil size may be an accessible means to address this research question.

The autonomic nervous system regulates pupil size through the control of two muscles in the pupil, the iris sphincter muscle which causes the constriction of the pupil and the dilatory muscle which promotes the dilation of the pupil (Larsen & Waters, 2018). Pupil size is dependent on the sympathovagal balance, with parasympathetic activity promoting pupil constriction through recruitment of the iris sphincter muscle via the midbrain Edinger-Westphal nucleus. Pupil dilation is dependent on the sympathetic system which inhibits the activity of the Edinger-Westphal nuclei leading to pupil dilation (Larsen & Waters, 2018). There is evidence to suggest that fluctuations in pupil size reflect an indirect readout of the changes in brain arousal during cognitive activity. Specifically, changes in pupil diameter have been hypothesised to be a readout of the activity of the noradrenergic neurons of the LC (Costa & Rudebeck, 2016). Evidence for the link between the LC and pupil size comes from the observation that the neuronal activity of the LC fluctuates almost simultaneously with changes in pupil diameter (Aston-Jones & Cohen, 2005; Nassar et al., 2012) and that the LC activity relates to pupil size, even in the absence of a cognitive task (Joshi et al., 2016). The activity of the LC and the diameter of the pupil were also correlated during a decision-making task (Varazzani et al., 2015). Furthermore, human studies combining fMRI and pupillometry have found activations in the area of the brainstem compatible with the LC were linked to fluctuations in pupil diameter, during resting state and for a novelty detection task (de Gee et al., 2017; Murphy et al., 2014).

Changes in pupil diameter can also be induced in response to cognitive effort which can be triggered by external stimuli (Joshi & Gold, 2020; Mathôt, 2018). In response to an external task event, the pupil dilates and then constricts back to baseline. This pupil response to a task event is called the ‘task-evoked pupil response’ (TEPR). These TEPRs can also be influenced by factors such as the demand of the cognitive task and the performance (Aston-Jones & Cohen, 2005; Kahneman & Beatty, 1966). An unestablished projection from the LC and the Edinger-Westphal nuclei has been hypothesised to drive TEPRs (Costa & Rudebeck, 2016; Smith et al., 2006), but the exact mechanism of the link between the size of the pupil and the activity of the LC is still not known. Studying TEPRs is nevertheless often considered a non-invasive means to determine the ongoing alterations in the LC phasic activity or arousal level during cognitive tasks.

The pupil is well known to adapt to changes in the light environment, with the pupil constricting at higher light levels (Larsen & Waters, 2018). However, whether the TEPRs are influenced by light’s NIF effects is currently not known. We, therefore, decided to study the TEPRs under different light conditions. We measured pupil diameter during two cognitive tasks and examined the effect of light level, expressed in melanopic (mel) Equivalent Daytime Illuminance (EDI) lux, on the TEPRs to auditory stimuli. We hypothesised that the TEPRs would be greater under higher irradiance levels due to the stimulating NIF impact of light. To test this hypothesis, we used eye tracking data from healthy young participants, who completed an attentional and an emotional auditory cognitive task during a functional Magnetic Resonance Imaging (fMRI) recording whilst exposed to different light conditions.

## Methods

### Participants

20 healthy participants (24.05 y± 4.0; 14 women) gave their written informed consent to take part in the study, which was approved by the Ethics Committee of the Faculty of Medicine of the University of Liège. The participants were assessed for the exclusion criteria with a semi-structured interview and questionnaires. None of the participants had a history of psychiatric and neurological disorders, sleep disorders, the use of psychoactive drugs or addiction. Participants had no history of night shift work during the last year or recent transmeridian travel during the last 2 months; excessive caffeine (>4 caffeine units/day) or alcohol consumption (>14 alcohol units/week); and were not taking medication or smoking. Their scores on the 21-item Beck Anxiety Inventory (Beck et al., 1988) and the Beck Depression Inventory-II (Beck et al., 1961) were minimal or mild (< 17) and minimal (< 14), respectively normal. Women were not pregnant or breastfeeding. Participants reported no history of ophthalmic disorders or auditory impairments and were screened for colour blindness. Due to technical issues (see below), 15 and 14 participants were, respectively, included in the analyses of the attentional and emotional tasks (**Table 1**).

**Table 1:**
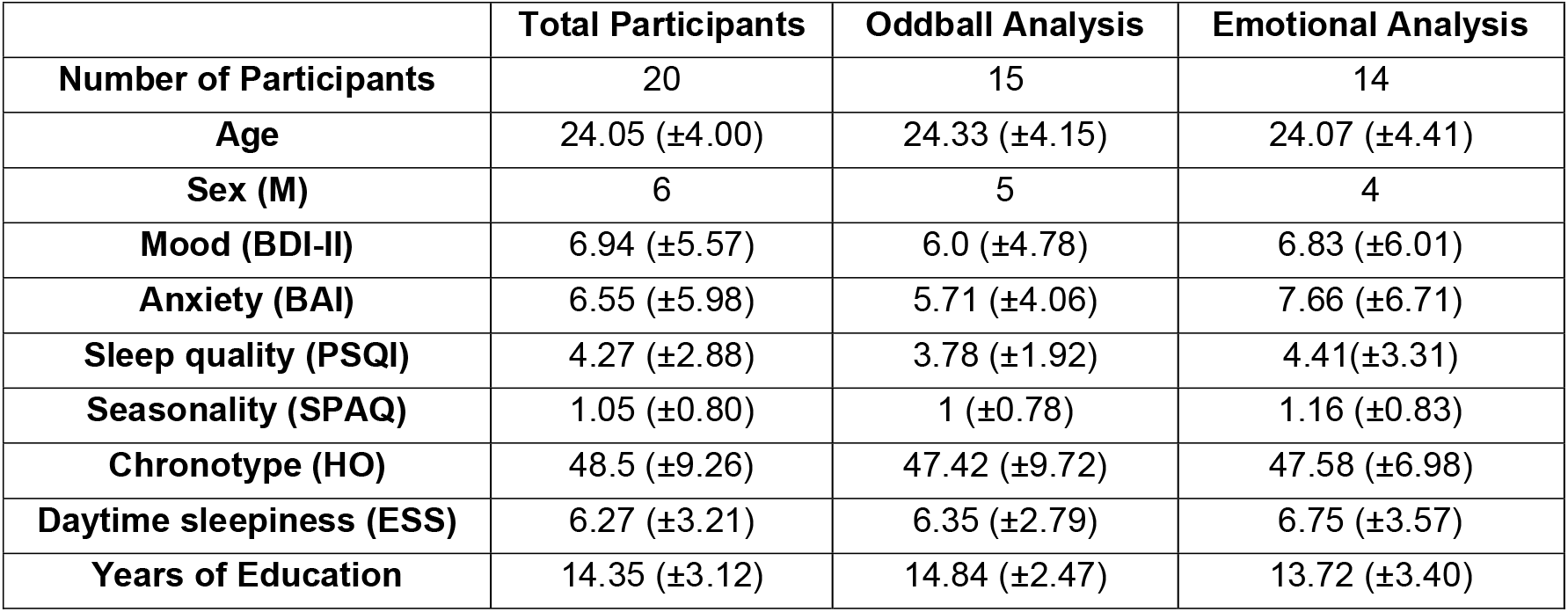
Table of participants included in the analysis. Columns for total number of participants who completed the study, and the number of participants included for each task. BDI-II, Beck’s Depression Inventory; BAI, Beck Anxiety Inventory; PSQI, Pittsburgh Sleep Quality Index; SPAQ, Seasonal Pattern Assessment Questionnaire; HO, Horne and Östberg; ESS, Epworth Sleepiness Scale. Refer to the main text for references.

|  | **Total Participants** | **Oddball Analysis** | **Emotional Analysis** |
| --- | --- | --- | --- |
| **Number of Participants** | 20 | 15 | 14 |
| **Age** | 24.05 ($\pm 4.00$) | 24.33 ($\pm 4.15$) | 24.07 ($\pm 4.41$) |
| **Sex (M)** | 6 | 5 | 4 |
| **Mood (BDI-II)** | 6.94 ($\pm 5.57$) | 6.0 ($\pm 4.78$) | 6.83 ($\pm 6.01$) |
| **Anxiety (BAI)** | 6.55 ($\pm 5.98$) | 5.71 ($\pm 4.06$) | 7.66 ($\pm 6.71$) |
| **Sleep quality (PSQI)** | 4.27 ($\pm 2.88$) | 3.78 ($\pm 1.92$) | 4.41 ($\pm 3.31$) |
| **Seasonality (SPAQ)** | 1.05 ($\pm 0.80$) | 1 ($\pm 0.78$) | 1.16 ($\pm 0.83$) |
| **Chronotype (HO)** | 48.5 ($\pm 9.26$) | 47.42 ($\pm 9.72$) | 47.58 ($\pm 6.98$) |
| **Daytime sleepiness (ESS)** | 6.27 ($\pm 3.21$) | 6.35 ($\pm 2.79$) | 6.75 ($\pm 3.57$) |
| **Years of Education** | 14.35 ($\pm 3.12$) | 14.84 ($\pm 2.47$) | 13.72 ($\pm 3.40$) |

Participants followed a loose sleep schedule (±1h from habitual bed/wake-up time) during the 7 days preceding the laboratory experiment to warrant similar circadian entrainment across participants (verified using wrist actigraphy and sleep diaries). They were asked to refrain from caffeinated and alcohol-containing beverages and excessive exercise for at least 3 days before the experiment. Participants were familiarised with the MRI environment one week before the experiment during an MRI session where structural images of the brain were acquired.

### Experimental Protocol

Most participants (N=17) arrived at the laboratory 1.5 to 2 hours after habitual wake time, while a minority (N=3) were admitted to the lab 1.5 to 2 hours before habitual bedtime. The study is meant to assess the time-of-day effect in the future. All results presented here consider time-of-day difference (see statistics) and all statistical output remains the same if we only include the data of the morning participants (data not shown).

Participants were first exposed for 5 minutes to a bright white light (1000 lux) and were then maintained in dim light (<10 lux) for 45 minutes to standardise participant light history before the fMRI session (**Fig.1A**). During this period participants were given instructions about the fMRI cognitive tasks and completed practice tasks. The fMRI session consisted of participants completing an executive task (25-min), an attentional task (15-min) and an emotional task (20-min) (**Fig.1B, C**). The executive task was always done first and then the order of the following two tasks was counterbalanced. Only the latter two tasks are discussed in the present paper as they consisted of streams of events, putatively triggering TEPRs (rather than blocks as in the executive task).

**Figure 1.**
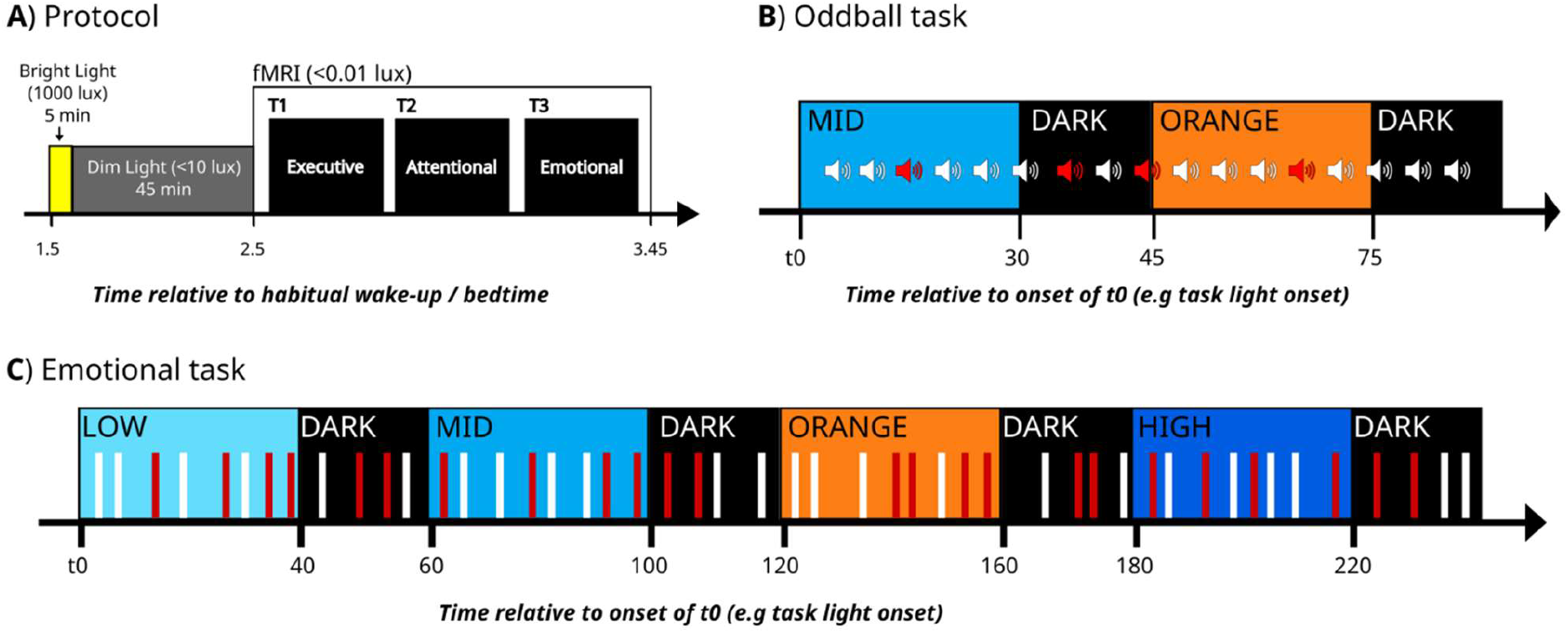
Experimental design. (**A**) General protocol. Time relative to scheduled wake-up/bedtime (hours). Following standardization of immediate prior light exposure (see methods), participants performed an executive (not discussed in the present paper), an attentional and emotional task in functional magnetic resonance imaging (fMRI). (**B**) Detailed procedures of the attentional task (oddball). Time (seconds) relative to t0, a time point arbitrarily chosen as the light onset of the session. The task consisted of a stream of standard sounds (80%) and pseudo-randomly interspersed odd sounds (20%), participants were asked to identify the odd stimuli through a button press. Whilst completing the task participants were exposed to white BEL (92 mel EDI lux; 6500K) (MID) and a monochromatic orange (0.16 mel EDI lux; 589mn) light. Light exposures lasted 30 sec and were separated by 10 sec periods of darkness. Odd (red) and standard (white) stimuli were equally distributed across the two light conditions and darkness. (**C**) Detailed procedures of the emotional task. Time (seconds) relative to t0, a time point arbitrarily chosen as the light onset of the session. The task consisted of a lure gender discrimination of auditory vocalizations of the three pseudo-word types (“goster,” “niuvenci,” or “figotleich”) while exposed to the alternating white BEL of three different intensities (37, 92, 190 mel EDI lux; 6500K) (LOW, MID, HIGH) and a monochromatic orange (0.16 mel EDI lux; 589mn) light. Light exposures lasted 30 to 40 sec and were separated by 15–20-sec periods of darkness. Untold to the participants, vocalizations were pronounced with angry (red bars) and neutral (white bars) prosody pseudo-randomly and equally distributed across the three light conditions.

An MRI-compatible light system designed-in-lab was developed to ensure relatively uniform and indirect illumination of participants’ eyes whilst in the MRI scanner. An 8-m long MRI-compatible dual-branched optic fibre (Setra Systems, MA, USA) transmitted light from a light box (SugarCUBE, Ushio America, CA, USA), that was stored in the MRI control room. The dual end of the optic fibre was attached to a light stand fitted at the back of the MRI coil. This allowed for equal illumination of the participants’ eyes. A filter wheel (Spectral Products, AB300, NM, USA) and optical fibre filters (a monochromatic orange light filter (589mn; full width at half maximum: 10 nm) and a UV long bypass (433 – 1650nm) filter) were used to create the light conditions needed for the experiment.

Both tasks were programmed with Opensesame (3.2.8) (Mathôt et al., 2012) and launched from a computer in the MRI control room. Participants heard the auditory stimuli through MR-compatible headphones (Sensimetrics, Malden, MA) and the volume was set by the participant before starting the tasks to ensure a good auditory perception of all the task stimuli. Participants used an MRI-compatible button box to respond to task items (Current Designs, Philadelphia, PA). During the attentional task, participants were exposed to 30s of light blocks separated by 10s of darkness (<0.1 lux). The light conditions used were a polychromatic, blue-enriched white light emitting diode (LED) light (92 mel EDI lux; 6500K) and a monochromatic orange light (0.16 mel EDI lux). The light blocks were repeated 7 times for each light condition. During the emotional task, participants were exposed to 30 to 40s periods of light blocks separated by 20s of darkness (<0.1 lux). The light conditions used were three different irradiances of a polychromatic, blue-enriched white LED light (37, 92, 190 mel EDI lux; 6500K) and a monochromatic orange light (0.16 mel EDI lux) (**Fig.2; Table 2**). The light blocks were repeated five times for each light condition.

**Table 2:** Light Characteristics. Additional light characteristics of the two light sources used. Blue enriched light (BEL) (low, mid, and high) and monochromatic light (589nm).

|  | **Low BEL** | **Mid BEL** | **High BEL** | **Monochromatic light (589nm)** |
| --- | --- | --- | --- | --- |
| **Lux** | 47 | 116 | 240 | 7.5 |
| **Peak Spectral Irradiance (nm)** | 460 | 460 | 460 | 590 |
| **Melanopic EDI (ipRGCs)** | 37 | 92 | 190 | 0.16 |
| **Rhodopic EDI (Rods)** | 39 | 97 | 201 | 0.94 |
| **Cyanopic EDI (S-cones)** | 32 | 79 | 163 | 0 |
| **Chloropic EDI (M-cones)** | 44 | 110 | 227 | 5 |
| **Erythropic EDI (L-cones)** | 46 | 113 | 233 | 8 |
| **Irradiance ($\mu\text{W}/\text{cm}^2$)** | 15 | 36 | 75 | 1.4 |
| **Photon flux($1/\text{cm}^2/\text{s}$)** | 4.12E+13 | 1.02E+14 | 2.10E+14 | 4.24E+12 |
| **Log Photon Flux ($\log_{10}$ ($1/\text{cm}^2/\text{s}$))** | 13.61 | 14.01 | 14.32 | 12.63 |
| **Narrowband peak** | - | - | - | 589 |
| **Narrowband FWHM** | - | - | - | 10 |

**Figure 2.**
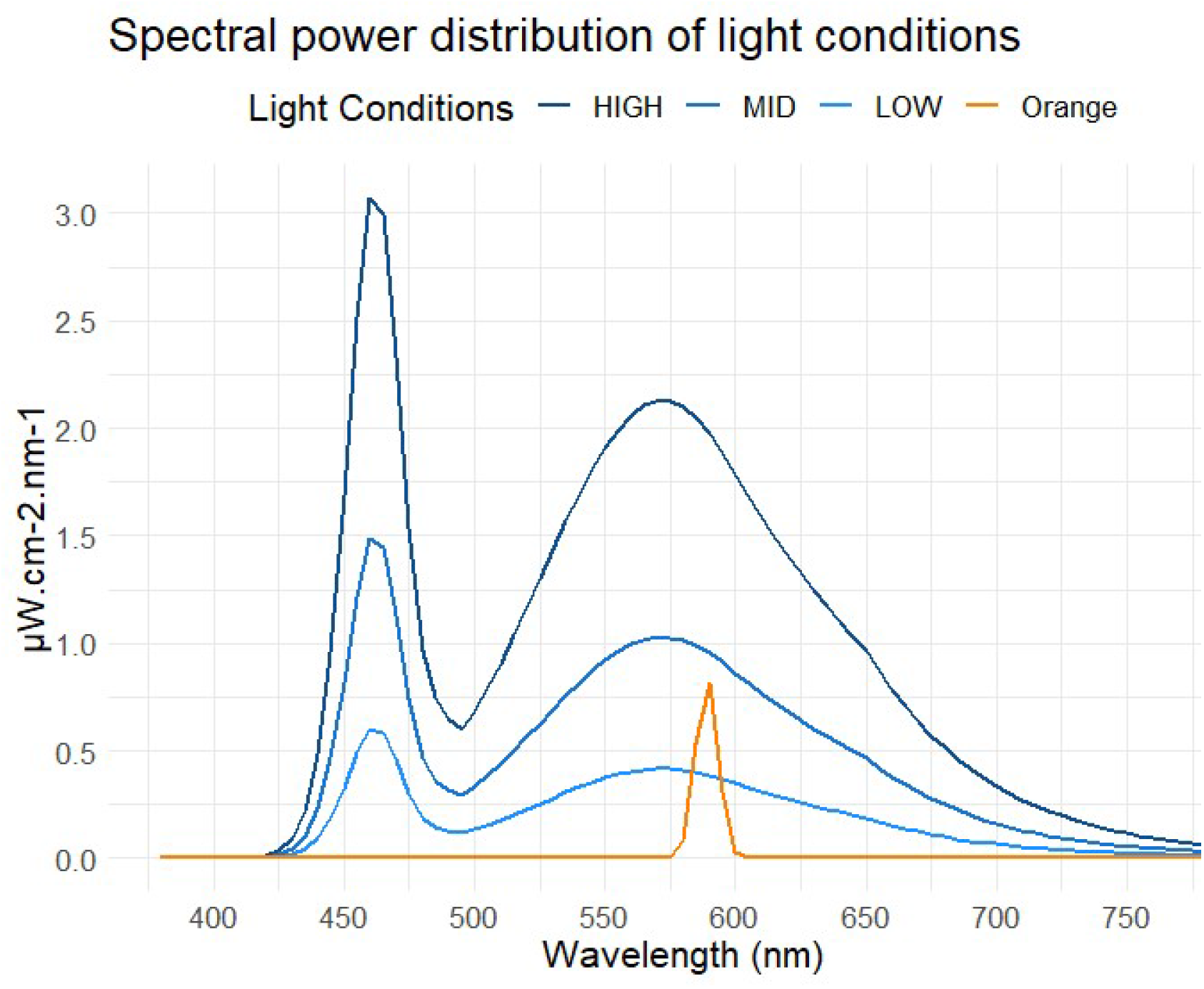
Spectral power distribution of light conditions. Orange: monochromatic orange light, 0.16 mel EDI lux, 589mn; BEL LOW, MID, and HIGH: light of three different intensities (37, 92, 190 mel EDI lux; 6500K). See table 2 for additional characteristics.

### Attentional Task

The attentional task used was a mismatch negativity or oddball task (Kiehl & Liddle, 2003). Participants were asked to detect a rare randomly occurring target (or odd) item in a stream of frequent standard items. They used the keypad to report the detection of the odd items. Stimuli (n=315) consisted of frequent standard ((500Hz, 100ms) and odd tones (1000 Hz, 100ms), presented 80% and 20% of the time, respectively, in a pseudo-randomized order. The inter-stimulus interval between stimuli was 2s. Target and standard stimuli were equally distributed across the two light conditions and the separating darkness periods (**Fig.1B**).

### Emotional Task

The emotional task used was a gender discrimination of auditory vocalizations task (Banse & Scherer, 1996). Participants were asked to use the keypad to indicate what they believed the gender of the person pronouncing each token was. The gender classification was a lure task ensuring participants paid attention to the auditory stimulation. The purpose of the task was to trigger an emotional response as participants were not told that 50% of the stimuli were pronounced with angry prosodies. The 240 auditory stimuli were pronounced by professional actors (50% women) and consisted of three meaningless words (“*goster*”, “*niuvenci*”, “*figotleich*”). The stimuli were expressed in either an angry or neutral prosody, which has been validated by behavioural assessments (Banse & Scherer, 1996)(Banse & Scherer, 1996)(Banse & Scherer, 1996) and in previous experiments (Grandjean et al., 2005; Sander et al., 2005). The stimuli were also matched for the duration (750 ms) and mean acoustic energy to avoid loudness effects. During each 30 to 40-s light block, four angry prosody stimuli and four neutral prosody stimuli were presented in a pseudorandom order and delivered every 3 to 5 seconds. A total of 160 distinct voice stimuli (50% angry; 50% neutral) were distributed across the four light conditions. The darkness period separating each light block contained two angry and two neutral stimuli. A total of 80 distinct voice stimuli (50% angry; 50% neutral) were distributed across the darkness periods (**Fig.1C**).

### Pupil

The right eye movements and the pupillary size was recorded continuously with an infrared eye tracking system (Eyelink-1000, SR Research, Osgoode, ON, Canada) (Sampling rate, 1000hz). Pupil data was analysed using MATLAB R2019b (MathWorks, MA, USA). Participants with more than 25% missing or corrupted eye-tracking data were excluded. Blink events were replaced with linear interpolation and the data was smoothed using the rlowess a robust linear regression function. The transient pupil response was computed as the change in the pupil diameter from before (baseline) and after (maximum) the auditory stimulus presentation. Baseline pupil diameter was computed as the mean pupil diameter over 1s before stimuli onset. The maximum pupil diameter was sought over a 1s window after sound onset. TEPRs were computed as the ratio between maximum and baseline diameter. For the attentional task, one participant was excluded because they did not complete the entire attentional task and four were excluded as there was over 25% missing or corrupt pupil data. Therefore, we included 15 participants in the analysis of the oddball task (Table 1). For the emotional task, two participants were excluded as there was over 25% missing or corrupt pupil data. One participant was excluded because he did not complete the entire emotional task correctly and three were excluded due to problems with the eye-tracking system. Therefore, we included 14 participants in the analysis of the emotional task (**Table 1**).

### Statistical analyses

Statistical analyses were computed using SAS 9.4 (SAS Institute, NC, USA) using individual TEPRs segregated per stimulus type and light condition. Values were considered outliers if they were >± 3 standard deviations (SD) across the entire dataset and were therefore removed. Analyses consisted of Generalised Linear Mixed Models (GLMM) seeking effects of light condition (i.e., mel EDI lux level) on the TEPRs. TEPRs were set as the dependent variable, with subject as a random factor, and light condition and stimulus type as repeated measures (autoregressive (1) correlation), together with the time of day, age, BMI and sex as covariates. GLMM were adjusted for the dependent variable distribution. Post hoc contrasts were corrected for multiple comparisons using a Tukey adjustment.

## Results

The performance of both tasks was high, with 96.6% (± 0.5) [mean (± SD)] of detection of target sounds during the attentional (oddball) task and 93.9% (±7.21) button response during the emotional (gender classification) task. In line with the literature (Sander et al., 2005; Vandewalle et al., 2010), for the emotional task, reaction times (RTs) were faster for neutral stimuli with a 1192ms (±182.8) average compared to 1234ms (±199.8) RT for emotional prosody vocal stimulation (*p*=0.0004) suggesting that the task was successful in triggering a differential response according to the emotional content. As no response was collected for the standard tone in the oddball task, RT could not be compared between stimulus types for the attentional task. Although not relevant to the task and not compromising any emotional effect (Grandjean et al., 2005), gender detection accuracy for the emotional task was relatively low - 79% ±11%.

It is well established that pupil size changes in response to variations in environmental irradiance. In a joint paper (Beckers et al., n.d.) (*BioRxiv*), we notably confirm this and report that the sustained constriction of the pupil increased with higher light levels in the same sample of participants that completed the same protocol. In contrast to the joint paper (Beckers et al., n.d.) (*BioRxiv*), here, we consider whether changes in light conditions, as indexed by mel EDI lux, impact the TEPRs for an attentional and emotional task. Both tasks consist of streams of events that putatively trigger TEPRs, and both have two types of auditory stimulations. We hypothesised that the TEPRs would be greater under higher light levels due to the stimulating NIF impact of light.

To characterise the effect of light conditions on TEPRs for the attentional task, an initial GLMM was conducted with TEPRs during the oddball task as the dependent variable. The results yielded significant main effects of stimulus type (target and standard tones; F_(1,1548)_ =189.27, *p*≤.0001) and light condition (F_(2,1548)=_ 13.71, p≤.0001). Importantly, the GLMM detected a significant interaction between stimulus type and light condition (F_(2,1548)_ =3.65, *p*=.02) (**Fig. 3A**). Post-hoc analyses first indicated that TEPRs were larger for target vs. standard stimuli (*p*≤.0001). They further indicated that TEPRs were smaller during darkness as compared to the blue-enriched light condition (92 mel EDI lux; *p*≤.0001) but not when compared to the orange (0.16 mel EDI lux; *p*=.1) light condition. However, TEPRs were significantly different when comparing the orange (0.16 mel EDI lux) light to the blue-enriched light (92 mel EDI lux; *p*=.002). Finally, post hoc analyses indicated that TEPRs significantly increased with higher light irradiance for the standard (*p*≤.0001) but not the target stimuli (*p*>.2).

**Figure 3.**
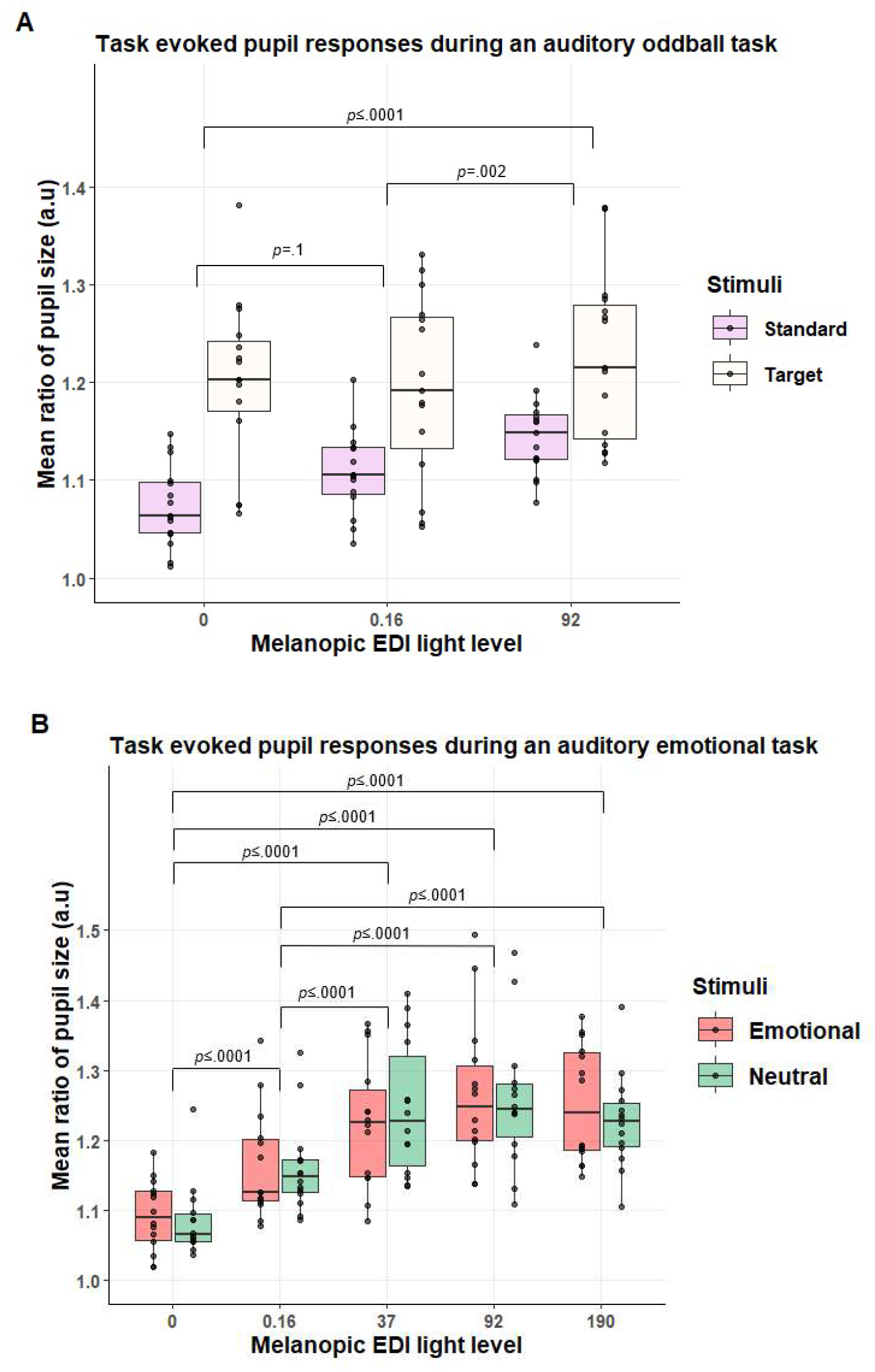
Task-evoked pupil response (TEPRs) across light conditions and stimulus type. **A**) TEPRs under the different light conditions during the attentional (oddball) task (N = 15; 24.33y (±4.15); 10 women). Individual average TEPRs were computed per stimulus type and light condition. TEPRs were significantly higher for target vs. standard stimulations (p=<.0001), as well as under higher vs. lower melanopic EDI light levels (p=<.0001). A significant light condition by stimulus type was also found (p=0.02) and post hoc analyses indicated that TEPRs significantly increased with higher light irradiance for the standard but not the target stimulations. **B**) TEPRs under different light levels during the emotional task (N = 14; 24.07y (±4.41); 10 women). Individual average TEPRs were computed per stimulus type and light condition. There was no significant difference between neutral and emotional stimulations (p=.8) while TEPRs were greater under higher vs. lower melanopic light levels (p=<.0001). There was no light condition by stimulus type interaction (p=0.7).

The second GLMM, with TEPRs during the emotional task as the dependent variable, led to a significant main effect of light condition (F_(4,1072)_= 77.78, p=≤.0001). Despite there being a qualitative difference between angry and neutral stimuli, there was no significant main effect of stimulus type (F_(1,1072)_ =.06, *p*=.8) (**Fig.3B**). In addition, there was no interaction between stimulus type and light condition (F_(4,1073)_ =.4, *p*=.7). Post hoc analysis showed a significant difference between darkness and all four light conditions (*p*≤.0001) as well as between the orange (0.16 mel EDI lux) light and the blue-enriched light conditions (37, 92, 190 mel EDI lux) (*p*≤.0001). There was no significant difference between the blue-enriched light conditions (37, 92, 190 mel EDI lux; p ≥.6).

## Discussion

TEPRs consist of transient pupil dilations triggered by the processing of stimulations over diverse cognitive domains. They are considered to be at least in part driven by a transient increase in the activity of the LC-NE system (Larsen & Waters, 2018). Here, we tested whether the TEPRs evoked by auditory stimulus during two cognitive tasks would be larger under higher ambient light levels to investigate whether light’s NIF impacts on cognitive brain activity could potentially be mediated through the LC. To test this hypothesis, we analysed eye tracking data from young healthy participants who completed an attentional and an emotional cognitive task during an fMRI protocol whilst exposed to different light conditions. The results reveal that despite a smaller sustained pupil size at higher light levels (Beckers et al., n.d.) (*BioRxiv*), the TEPRs to auditory stimulus were larger under higher light irradiances, as indexed by mel EDI lux. Although this main finding was detected for both the attentional and emotional tasks, we further observed task-specific differences in the impact light irradiance has on the different types of stimuli of each task.

The LC is involved in the processing of salient events through an increase in its phasic activity (Berridge & Waterhouse, 2003). The oddball task, which mimics novelty/salience detection has been previously used to assess the phasic activity of the LC (Rajkowski et al., 1994), while the LC is also known to be important for emotional processing (Aston-Jones & Cohen, 2005; Bradley et al., 2008). The oddball task was reported to trigger TEPRs (both in the visual and auditory modality) that were larger for the odd target stimuli, which is in line with our findings (Gilzenrat et al., 2010; Murphy et al., 2014). Similarly, TEPRs were also reported using emotional tasks (Aston-Jones & Cohen, 2005; Bradley et al., 2008). Pupil size depends on the parasympathetic-sympathetic balance and transient pupil dilation is thought to reflect an increase in arousal due to an increase in the sympathetic tone (Larsen & Waters, 2018). Although recent investigations indicated that it is likely not the sole driver of transient pupil dilation, *in vivo* animal studies support that transient increases in pupil size were directly related to the firing of the neurons of the LC (Costa & Rudebeck, 2016).

Our findings suggest therefore that the phasic activity of the LC related to an ongoing cognitive process is likely to be affected by changes in ambient light level. The LC is a good candidate to mediate the impact of light on human alertness and cognition through an effect on other subcortical and cortical structures (Aston-Jones & Cohen, 2005). The thalamus pulvinar could likely be one of these downstream structures as it is the most consistently affected by light in previous investigations on the impact of light on non-visual cognitive brain activity (Vandewalle et al., 2009). Other structures and nuclei, e.g., within the hypothalamus or basal forebrain, could also be implicated, while the recruitment of limbic and cortical areas would depend on the ongoing cognitive processes (Gaggioni et al., 2014).

Our results indicate that the impact of increasing light level is stronger for standard compared with target stimulation. We interpret this as a ceiling effect for TEPRs elicited by target stimulations that cannot be further increased, while the milder TEPRs triggered by standard stimulations in darkness or at lower light levels can be increased under higher ambient light. In line with this interpretation, the impact of light on non-visual cognitive brain activity was previously found to be reduced in the evening, when the endogenous circadian signal promoting wakefulness is strongest and therefore when alertness could not be further increased (Vandewalle et al., 2011). In contrast, lights’ impact was increased in the morning following sleep deprivation, when the circadian signal is weaker and the need for sleep is high, and therefore when alertness can benefit from the external stimulating impact of light (Vandewalle et al., 2011). If our interpretation is correct, this could mean that light can only affect the activity of the LC when it is not already highly recruited by the processing of a salient stimulation.

The situation is different if we consider the emotional task as we find no difference between the TEPRs triggered by the emotional and neutral simulations. This could call into question the emotional valence of the stimuli included in the task. However, the emotional task has been previously extensively validated and was successful in triggering differential brain responses to emotional vs. neutral stimulations, including in studies interested in the NIF effects of light (Banse & Scherer, 1996; Grandjean et al., 2005; Vandewalle et al., 2010). We also find that RTs were significantly slower in response to emotional vs. neutral stimulations, which is in line with the literature and supports that the emotional valence of the stimuli was perceived by the participants (Sander et al., 2005; Vandewalle et al., 2010). Yet, the emotional response may not be strong and/or different enough from the response to neutral stimuli to be detected with 15 subjects. Auditory emotional stimuli are indeed considered to be less effective at provoking an emotional response when compared to visual emotional stimuli (Bradley et al., 2008). It may also be that the unexpected occurrence of neutral stimulations (stimulations were pseudo-randomly delivered every 3 to 5 seconds) triggers a TEPR that is similar to the emotional stimuli. Our results further indicate that given the relatively mild response elicited in darkness or at lower light levels, TEPRs could be increased by increasing light levels. The maximum increase seems to be reached already with the lower level of polychromatic, blue-enriched white light (37 mel EDI lux) to ceil thereafter. Interestingly, the maximum TEPRs for both the oddball and emotional tasks seem to lay on average around 1.25, i.e. a 25% increase on average in pupil size compared to baseline (cf. **Fig 2**).

We emphasize that our study has limitations. The light conditions included do not allow for determining which of the human photoreceptors are mostly contributing to the TEPRs. Rods, cones and ipRGCs could equally be involved with differential recruitment at the different light levels we used (Mure, 2021). Future research could use metameric light sources with which the wavelength compositions can be manipulated to differentially recruit one photoreceptor type while leaving the others relatively similarly recruited (Viénot et al., 2012). We also stress that we did not have access to the brain activity associated with TEPRs. The assumptions made regarding the recruitment of the LC can only be verified using the fMRI data acquired simultaneously with the pupil data.

## Conclusion

Overall, this study shows that two seemingly opposite NIF impacts of light can be detected simultaneously when focusing on pupil size with concomitant sustained pupil constriction and transient pupil dilation induced by increasing light levels. This is true for two different auditory cognitive tasks while increased transient pupil dilation may only be possible if TEPRs are not already at maximum. Given the putative link between LC phasic activity and transient pupil dilation (Costa & Rudebeck, 2016), the results presented here provide further support for the involvement of the LC in the stimulating impact of light on alertness and cognition.

## Abbreviations

EDI: Equivalent Daylight Illuminance
fMRI: Functional Magnetic Resonance Imaging
GLMMs: Generalised Linear Mixed Models
ipRGCs: Intrinsically Photosensitive Retinal Ganglion Cells
LC: Locus Coeruleus
LEDs: Light Emitting Diodes
mel: Melanopic
NIF: Non-Image-Forming
RTs: Reaction Times
SD: Standard Deviation
TEPR: Task-Evoked Pupil Response

## Acknowledgement

The authors thank, Christine Bastin, Annick Claes, Fabienne Collette, Christian Degueldre, Catherine Hagelstein, Gregory Hammad, Brigitte Herbillon, Patrick Hawotte, Sophie Laloux, Erik Lambot, Benjamin Lauricella, André Luxen, Christophe Phillips, Pierre Maquet, and Eric Salmon for their help over the different steps of the study.

This study was supported by the Belgian Fonds National de la Recherche Scientifique (FNRS; CDR J.0222.20), the European Union’s Horizon 2020 research and innovation program under the Marie Skłodowska-Curie grant agreement No 860613, the Fondation Léon Frédéricq, ULiège - U. Maastricht Imaging Valley, ULiège-Valeo Innovation Chair “Health and Well-Being in Transport” and Sanfran (LIGHT-CABIN), the European Regional Development Fund (Biomed-Hub), and Siemens. None of these funding sources had any impact on the design of the study nor on the interpretation of the findings. AB is supported by Synergia Medial SA and the Walloon Region (Industrial Doctorate Program, convention n°8193). EB is supported by the Maastricht University - Liège University Imaging Valley. RS and FB are supported by the European Union’s Horizon 2020 research and innovation program under the Marie Skłodowska-Curie grant agreement No 860613. IC, EK, IP, NM, GV are supported by the FRS-FNRS. SS was supported by ULiège-Valeo Innovation Chair and Siemens Healthineers.

